# Encoding and combining naturalistic motion cues in the ferret higher visual cortex

**DOI:** 10.1101/2025.09.23.678183

**Authors:** Thomas Schaffhauser, Pascal Mamassian, Yves Boubenec

## Abstract

Natural visual scenes contain rich flows of pattern motion that vary not only in orientation but also in spatial sizes and temporal rhythms. To properly interpret the motion, the brain must extract individual features and integrate them into coherent parts, and sometimes also segregate these parts from each other. Here, we performed single-neuron recordings in the motion-sensitive higher-order visual cortex (PMLS) of awake ferrets to investigate how complex motion signals are encoded and combined. We presented motion clouds—naturalistic stimuli with parametrically controlled spatiotemporal frequency content—and found that motion features were encoded in a temporally ordered sequence: orientation and spatial frequency emerged within 120 ms after stimulus onset, while temporal frequency and direction followed at later latencies. Time-resolved decoding revealed that this selectivity evolved dynamically within neurons and was distributed across the population. To probe motion integration, we introduced compound motion clouds composed of two or three localized frequency components. Neuronal responses were well explained by a linear pooling model, suggesting a simple summation mechanism of the individual components. However, a distinct subset of neurons exhibited late responses sensitive to changes in speed content despite matched marginals, consistent with receptive fields differentiating along the speed gradient. Together, we have uncovered a structured and distributed code for motion in high-level visual cortex, and provide mechanistic insights into how the brain parses complex motion in natural scenes.

## Introduction

Real-world motion is composed of multiple overlapping signals, each defined by distinct orientation, spatial frequency, and temporal frequency content. The visual system, stormed by this diversity of features, must reliably link local information, while accurately differentiating moving objects, and ultimately arbitrate between integration and segmentation of visual inputs (Braddick, 1993). Decades of work in humans and non-human primates have mapped the broad stages of this transformation from initial filtering in primary visual cortex (Movshon & Newsome, 1996) to integration in extrastriate areas such as MT (Middle Temporal, Albright, 1984; Born & Bradley, 2005), where selectivity for direction and speed emerges through combinations of spatial and temporal information (Simoncelli & Heeger, 1998). Each neuron contributes within this distributed population code to a pooled estimate of motion’s properties, enabling near-optimal readout of motion parameters (Salinas & Abbott, 1994; for a review, see Lisberger, 2010). This encoding-decoding cascade accounts for diverse visual behaviors, including smooth pursuit (Osborne et al., 2004), saccadic suppression and movement cancellation (Masson & Castet, 2002), pursuit of simultaneous targets (Lisberger & Ferrera, 1997), spatial pooling (Lorenceau & Zago, 1999), and motion transparency (Treue et al., 2000).

Despite detailed descriptions of visual motion processing under simplified laboratory conditions, how high-dimensional, naturalistic motion is encoded and decoded in the brain remains unclear. Natural scenes do not consist of single gratings or isolated motion vectors; instead, they contain concurrent, overlapping streams of motion (Braddick, 1993; Lorenceau, 2010). Encoding such stimuli involves multiple spatiotemporal frequency channels, and may engage distinct rules for combining or segregating motion components. Consistent with this proposal, early work using elementary stimuli like gratings or random dot motion suggested that neurons sum individual motion components (Treue et al., 2000; Zaharia et al., 2019), while processing of evidence from more complex natural scenes pointed to nonlinear interactions between speed channels (Inagaki et al., 2016; Gekas et al., 2017; Meso et al., 2022). In parallel, psychophysical studies have highlighted that human observers range from an *averaging* behavior across components (Lisberger & Ferrera, 1997; Lorenceau & Zago, 1999) to a *winner-take-all* behavior, depending on context (Busettini et al., 1996; Sheliga et al., 2006). Direct evidence to study this phenomenon from the activity of neuronal populations stimulated under naturalistic conditions has remained so far elusive.

Here, we address this challenge by recording single-neuron responses in awake ferrets to a novel class of stimuli called motion clouds, synthetic textures that approximate the statistics of natural motion while allowing precise control over their spatiotemporal frequency content (Leon et al., 2012; Vacher et al., 2018). We target area PMLS (Posterior-Medial Lateral Sulcus), a higher-order visual region homologous to primate MT (Lempel & Nielsen, 2019), to ask how motion features are encoded in time and combined across frequency channels. Our results reveal a structured temporal progression in feature encoding: orientation and spatial frequency emerged early, followed by temporal frequency and direction. We then show that multiple motion components are combine through a simple linear pooling rule, and identify a subpopulation of neurons selectively tuned to differences in speed content. These findings offer a mechanistic account of how higher-order cortex extracts, combines, and segregate motion information from complex inputs.

## Results

### Neuronal responses to naturalistic rich motion stimuli in ferret higher-order visual cortex

We recorded neuronal activity from the motion-processing area PMLS in awake ferrets to examine early sensory responses to wide-field naturalistic textures with rich spatio-temporal content, namely motion clouds (Figs 1a–c). Motion clouds (MCs) are a class of broadband white-noise stimuli: they differ in Fourier space from drifting gratings in that they are not confined to a single point but encompass a distribution of spatial and temporal frequencies. Their broadband nature resembles naturalistic motion while allowing a control over spatio-temporal frequency distribution (Leon et al., 2012; Vacher et al., 2018). In our experiments, MC had the mode of their spatial frequencies’ envelope ranging from 0.035 to 0.14 cyc/deg and temporal frequencies’ envelope ranging from 0.35 to 1.4 Hz across 8 directions (Fig 1d). The set of MCs covered five spatial frequencies, five temporal frequencies, and were aligned along five parallel speed lines, with, therefore, some pairs sharing similar marginal spatial frequency, temporal frequency, or speed power. They were viewed binocularly, and presented in a pseudo-random order for 500 ms, interleaved by a 250 ms luminance-matched uniform gray screen (Fig 1e).

**Fig 1.**
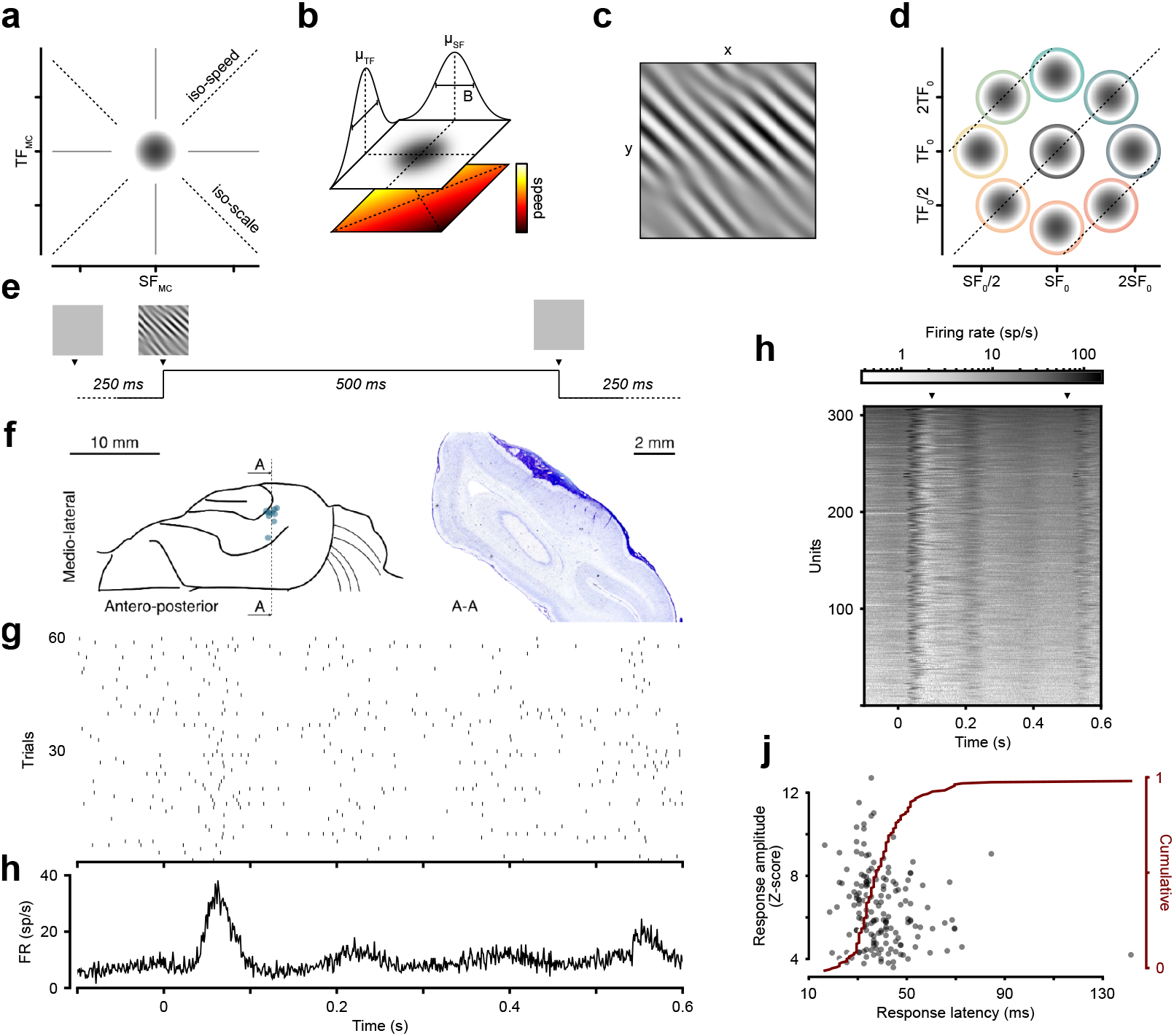
Motion clouds and responses in ferret higher-order visual cortex. **(a)** Presentation of the stimulus and its representation in log-log Fourier space where the x-axis is the spatial frequency (SF) and the y-axis the temporal frequency (TF). In this panel, MC stands for the critical frequencies of the motion cloud. **(b)** Representation of a motion cloud in the spatio-temporal log-log-space and its parametrization by spatial and temporal frequency modes and bandwidth (B = ^1^*/*_3_ of octave). Speed increases as the ratio of temporal over spatial frequencies. **(c)** Example frame of a motion cloud that was drifting to the top-right. **(d)** Positions of the nine MCs used in the experiments in log-log-space. In the first experiment, the central motion cloud had a SF_0_ of 0.07 cyc/deg and TF_0_ of 0.7 Hz, while in the second experiment, these values were shifted to 0.14 cyc/deg and 1.4 Hz while keeping the same 10 deg/s speed. **(e)** Sequence of stimulus presentation. A gray display was presented for 250 ms followed by one of the motion cloud stimuli for 500 ms. **(f)** Histological cut along the medio-ventral axis and stereotaxic location of the recordings. The blue dots show the locations of the recordings. **(g)** Raster plot for one example neuron and **(h)** its corresponding peristimulus time-histogram. Neurons had a fast and early transient response to stimulus onset, followed by a suppression for the rest of the stimulus duration, and a weaker offset response. **(i)** Collapsed evoked responses of all neurons analyzed in the first recording. Arrows indicate the beginning and end of the motion clouds presentation. Units were ranked by the overall strength of their response. **(j)** Distribution of responses onset latencies. Only the neurons with an excitatory response whose onset response reached at least 2 standard deviations above the baseline (plotted on the y-axis) were kept for this plot. The red curve shows the cumulative distribution.

A total of 530 units were recorded across two animals (experiment 1: 247 units for ferret A, and 62 units for ferret B; experiment 2: 177 units for ferret A, and 44 units for ferret B) in the right posterior-medial lateral sulcus (PMLS) (Fig 1f), a region involved in motion processing in the ferret (Philipp et al., 2005; Jarosiewicz et al., 2012; Lempel & Nielsen, 2019). A majority of units showed a fast transient response to the stimulus onset (median = 34 ms) followed by a suppression phase (example unit in Figs 1g, h, all units in Fig 1i). Response onset latencies were short, ranging from 18 to 68 ms (95% CI) post-stimulus onset, with a mean of 37.2 ± 1.0 ms (sem; Fig 1j), consistent with the fact that PMLS receives a direct input from the primary visual cortex in the ferret (Jarosiewicz et al., 2012) and overall low latencies generally observed in the ferret visual system (Alitto & Usrey, 2004).

### Sequential and distributed encoding of motion features

Neurons responded selectively to these dimensions, and this selectivity evolved over time. Most units displayed early orientation-tuned response (Fig 2a and Fig 2b, left), followed by direction tuning at later latencies (Fig 2a and Fig 2b, right), a response profile that suggests a temporal multiplexing of visual information. Neurons encoded different visual features of motion clouds with distinct temporal dynamics. Given that motion stimuli are characterized by a set of attributes with different complexity, we hypothesized that these features could be processed on different timescales. In particular, some attributes can be inferred from a single snapshot, like orientation and spatial frequency, while others require more processing time, like direction and temporal frequency, making the latter two attributes more complex than the former two. To test this hypothesis, we leveraged the parametrized nature of the motion clouds and applied cross-validated, time-resolved multi-class decoding to single-neuron activity for each feature. A majority of neurons (95%) showed significant decoding for at least one feature (Figs 2c–i), and 23% decoded all four features, namely orientation, spatial frequency, temporal frequency, and direction. Orientation showed better and earlier decodability than other attributes, rising sharply after stimulus onset and peaking around 120 ms (Fig 2d). Decoding of spatial frequency (Fig 2e), temporal frequency (Fig 2f) and direction (Fig 2g) followed, each reaching their peak at progressively later times (Figs 2h, i): first orientation (median = 117 ± 16 ms; 95% bootstrap CI), then spatial frequency (183 ± 25 ms), then temporal frequency (267 ± 67 ms), and finally direction (283 ± 84 ms). Feature decoding appeared distributed across the population, as units ranked by orientation decoding did not rank similarly for other features (Figs 2c–i). These results were stable across decoding windows’ sizes (Fig S2a), and confirmed by an integrative decoder (Fig S2b), and indicate that features selectivity built up with temporally staggered dynamics across neurons, consistent with a distributed and dynamic encoding scheme.

**Fig 2.**
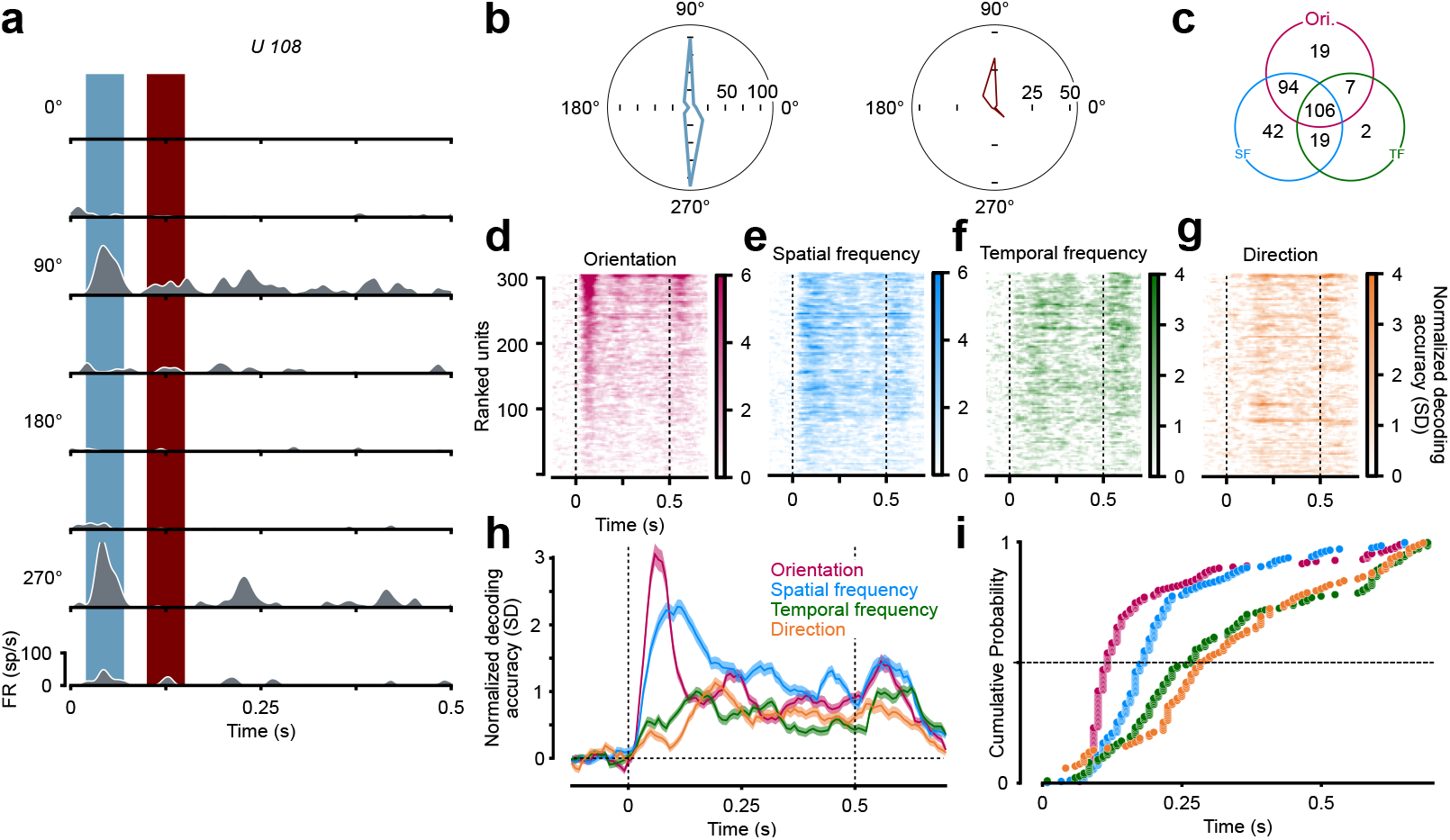
Build-up of visual feature selectivity. **(a)** Example of spiking activity for one unit in response to the 8 motion directions. The vertical regions indicate the early (blue) and late (red) windows used to estimate the tuning curve displayed in **b. (b)** Orientation tunings for the example cell in **a**. The panel on the left shows the orientation tuning during the early time window post-stimulus 20–70 ms, and the panel on the right the tuning for the late 100–150 ms time window. This cell is first tuned for orientation and then for direction. **(c)** Number of single unit successfully decoding for orientation, spatial, and temporal frequencies, or combinations of these stimulus features, out of a total number of 309 units analyzed. **(d)** Single unit decoding performance across time for orientation using a linear discriminant analysis (see *Materials and methods*). The 312 units recorded from two animals (247 for ferret A and 62 for ferret B) are ranked by their decoding performance. Vertical dashed lines indicate the onset and offset of the stimuli. **(e)** Decoding performance for spatial frequencies. Units are still ranked by their decoding performance on orientation for comparison. **(f)** Decoding performance for temporal frequencies, with units ranked by their decoding performance on orientation. **(g)** Decoding performance for directions, with units ranked by their decoding performance on orientation. **(h)** Normalized decoding performance for the four different features of the motion clouds, averaged across all neurons (colors as in **c–f**). **(i)** Cumulative probability of units response decoding peak time for each condition (colors as in **c–f**). This plot shows that half of the units (cross section of the plot at a height of 0.5) respond first to orientation, then spatial frequency, then temporal frequency, and finally direction.

### Response summation to concurrent stimuli

Although motion clouds share similarities with the spatio-temporal frequency content of natural visual scenes, these scenes often contain multiple and distinct components each with their own spatial and temporal frequency contents. To understand the mechanisms behind the combination of individual components, a set of pattern compound motion clouds (CMCs) were generated by combining two (or three) MCs. Such stimuli were constructed by summing individual components (Figs 3a, b) that were chosen to have different positions relative to the center of the spatio-temporal frequency space (SF_0_, TF_0_). To ensure matching contrast with the motion clouds, the CMCs were then normalized. The majority of the units responded to all the stimuli presented (56% reached a z-score above 2 for every stimuli presented).

**Fig 3.**
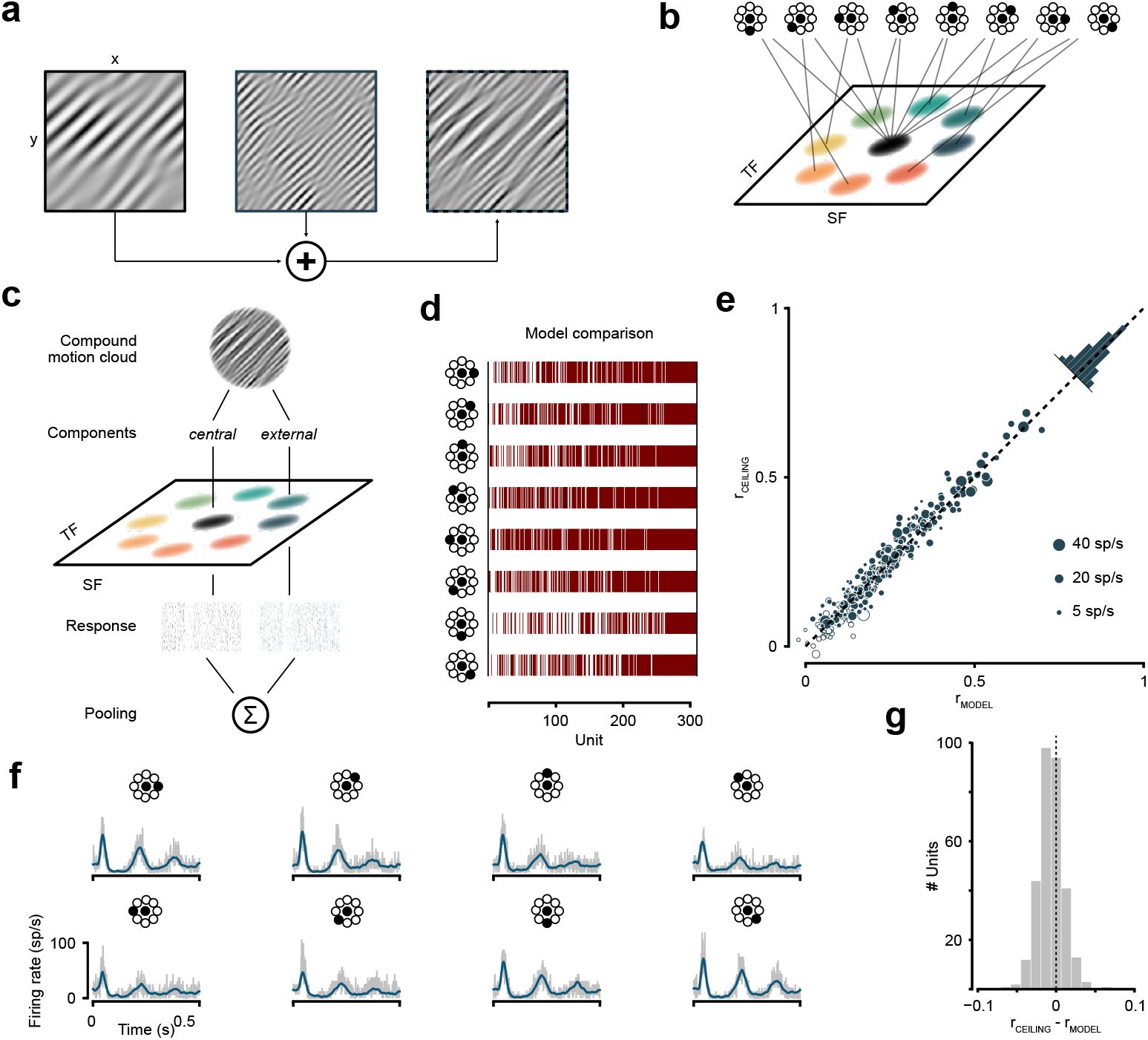
Compound motion clouds and summation of responses to concurrent stimuli. **(a)** Construction of the Compound Motion Clouds (CMCs) by superposition of multiple (here 2 are illustrated) motion clouds. **(b)** The different CMCs were designed by superposition of the central motion cloud to each of the 8 other motion clouds. **(c)** Model of response to a compound motion cloud (CMC) built from the combination of two motion clouds (the central MC is always part of all CMCs). **(d)** Model comparison across the eight CMCs. For each CMC, the summation, the full, and the winner-take-all models were tested and the best model was selected by the AIC measure. The winning model is shown by a line whose color is red if the summation model won, white if it is the winner-take-all model, and black the full model. Units are inversely ordered by the proportions of the eight CMCs that were overall best explained by the summation model. **(e)** Goodness of fit of the summation model. Firing rate predictability for the summation model is compared to the maximum achievable predictability obtained by test-retest reliability. This plot shows the analysis for one compound motion cloud. **(f)** Fit example for one unit with the firing rate for each compound motion cloud (in gray) and the instantaneous firing rate predicted by the model (in blue). **(g)** Difference between the prediction accuracy of the summation model (r_model_) and the test-retest correlation (r_ceiling_) for all conditions concatenated (for separated histograms for each compound motion clouds, see Fig S4a).

To characterize the response to compound motion clouds, we compared the performance of different models predicting unit responses to CMCs from the sum of the two MCs (*summation* model), their non-linear sum (*full* model, defined as a p-norm), or one of them (*winner-take-all* model) (Figs 3c, d; see *Materials and methods*). The *summation* model outperformed the *full* model and the *winner take-all* model in 73% of the cases (Fig 3d). At the population level, the model’s prediction varied by, at most, 5% from the linear prediction (see *Materials and methods*). This trend was observed across all CMCs presented and indicates that neurons responses to CMCs agree with a summation rule.

To further quantify the agreement between the summation model’s predictions and the responses to the CMCs, we measured the correlation between predictions and data in a cross-validated manner. Model predictions varied across neurons (Figs 3e, g; Fig 3f for an example fit), but these variations in fit quality (r_model_) were explained by differences in response reliability measured across repeats (r_ceiling_; Figs 3e, g; Pearson’s r ≥ 0.96; p < 0.001 for all conditions). This relationship held across the tested range of spatio-temporal combinations tested over all the pair CMCs where r_ceiling_ and r_model_ were not statistically different (|r_c_ − r_m_| ≤ 0.009; p ≥ 0.21 for all conditions; 1000 permutations; Fig S4a). These results confirm that neuronal responses closely approximated the vector average of their individual motion components and that variability in model fit reflected noise rather than deviations from summation. This principle extended to CMCs composed of three MCs (Figs S4b, c).

### A subset of neurons in PMLS have receptive fields tuned towards speed differences

To further probe how PMLS neurons process stimuli with rich content in the spatio-temporal domain, we performed an additional experiment with compound motion clouds composed of three motion clouds. These stimuli were built by combining MCs with different angles *φ* in Fourier log-log-space relative to the central motion cloud (Figs 4a, b). Altogether, there were four distinct triplet CMCs that were aligned along the iso-TF (*φ* = 0°), iso-speed (*φ* = 45°), iso-SF (*φ* = 90°) or iso-scale (*φ* = 135°) axes. In contrast to the pair CMCs (Figs 3a, b), these triplet CMCs had mean speeds, mean spatial and temporal frequencies that were identical to the central MC, and only differed in their frequency range along the different orientations *φ*. Notably, the CMCs that were aligned along the iso-speed and iso-scale axes were matched in spatial and temporal frequency content and varied only in their speed distribution.

**Fig 4.**
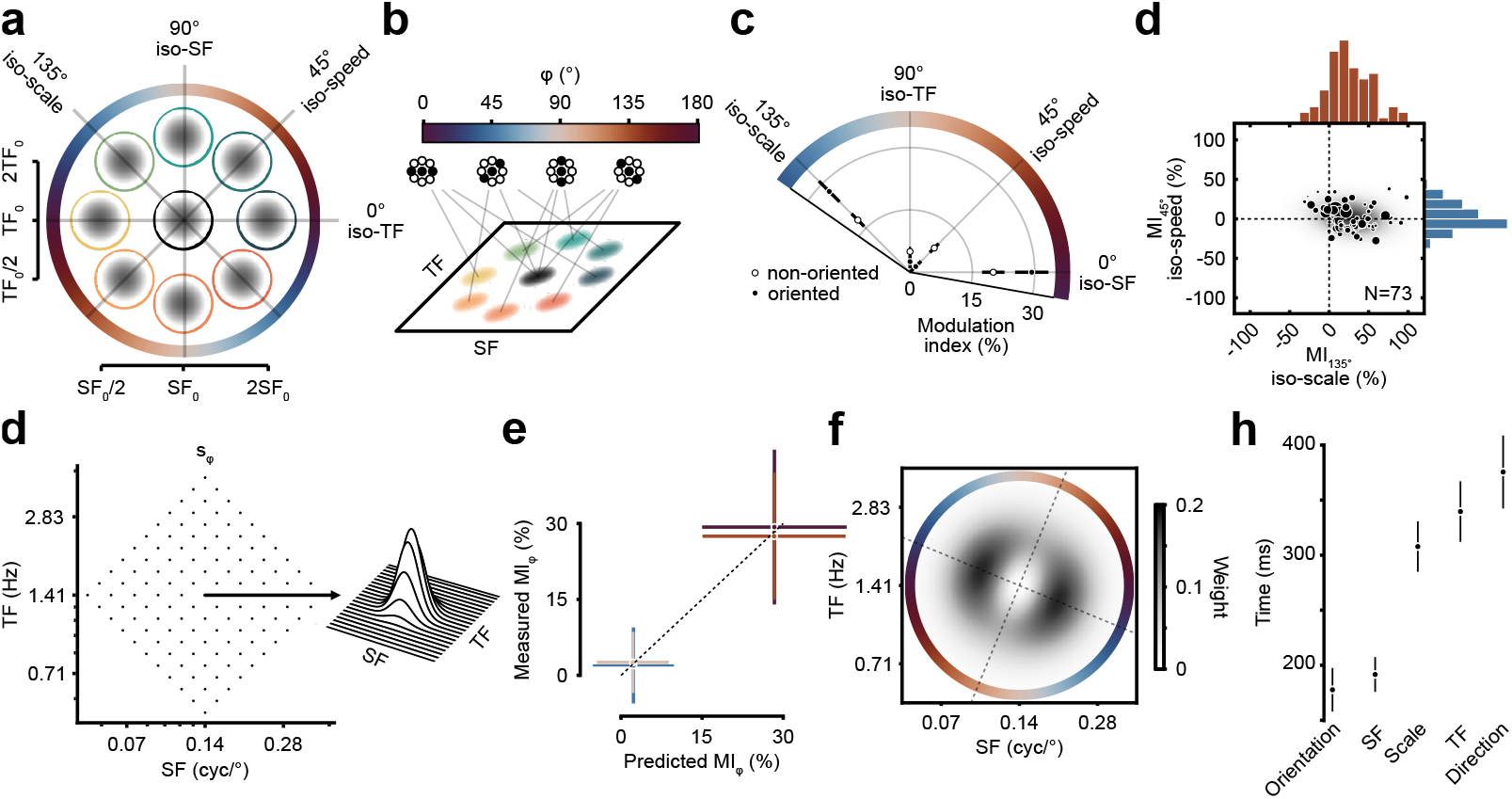
Responses to triplet compound motion clouds and receptive fields tuned for speed-base segmentation. **(a)** Representation of the different compound motion clouds built from a combination of three motion clouds. The central MC was always included, and two other MCs on either side were chosen along the iso-TF, iso-speed, iso-SF, or iso-scale axes. **(b)** Construction of the four compound motion clouds in Fourier space. The CMCs aligned along the iso-speed and iso-scale axes do not differ in their marginal spatial and temporal frequency energy, but are either consistent with a single speed (iso-speed) or multiple speeds (iso-scale). **(c)** Mean modulation indices (MI) for each CMC. The polar plot shows the MI for the units that decoded stimuli oriented in Fourier space (oriented units, filled circles) and units that failed to significantly decode stimuli oriented in Fourier space (non-oriented units, open circles). **(d)** Distribution of the modulation index along the iso-speed and iso-scale axes for the oriented units. **(e)** Illustration of the model for the response of PMLS neurons to the triplet CMCs. In this model, stimuli are analyzed by a bank of spatio-temporal frequency channels whose receptive fields are sensitive to the speed gradient. The response of these channels are then summed with different weights assigned to different spatio-temporal channels. **(f)** Predicted and measured modulation index for the four triplet CMCs. Error-bars represent bootstrap confidence intervals. **(g)** Fitted population receptive fields (weights *w*) in the spatio-temporal domain. Best fit parameters for the data obtained were are *σ*_*e*_[log_2_] = 0.43 ± 0.01, *σ*_SF_[log_2_] = 0.34 ± 0.02, *σ*_TF_[log_2_] = 0.37 ± 0.02, and *ρ* = 0.09 ± 0.01 (see *Materials and methods*) **(h)** Decoding onset latencies of the units that could significantly decode iso-speed from iso-scale stimuli, in comparison with latencies for orientation, spatial and temporal frequency, and direction.

Units were classified based on whether the stimulus could be decoded from their evoked response. A total of 73 units (33%) significantly discriminated between the iso-scale and iso-speed CMCs, these two CMCs having identical marginal spatio-temporal content. They were coined *“oriented”* units, in contrast to *“non-oriented”* units that lacked such selectivity. We further characterized their tuning properties by computing a modulation index MI_*φ*_ that quantified changes in response as stimulus’ frequency range increased along the axis *φ* (Figs 4c, d; see *Materials and methods* for details). Non-oriented units showed minimal modulation as the range in frequencies increased along the iso-temporal (*φ* = 0°; p = 0.08, 1000 permutations), iso-speed (*φ* = 45°; p = 0.29), iso-spatial (*φ* = 90°; p = 0.36) and iso-scale axes (*φ* = 135°; p = 0.08). In contrast, *oriented* units exhibited stronger responses when the frequency range increased along the iso-scale (+27 ± 4%; p = 0.04) and the iso-TF axes (Fig 4c, +29 ± 4%; p = 0.03). Notably, 85% of the oriented units preferred increasing content along the iso-scale axis (Fig 4d; MI_135°; iso-scale_) i.e. along the speed gradient, but not along the iso-speed axis (MI_45°; iso-speed_; p = 0.48).

This response profile diverges from predictions made for classic speed-tuned neurons, neurons whose receptive field are oriented along the iso-speed axis. In this scenario, firing rate is expected to increase when power expands along the iso-speed axis (MI_45°; iso-speed_ > 0), a prediction inconsistent with our data (Fig 4d, blue histogram; MI_45°; iso-speed_ ≈ 0). Alternatively, neurons showing speed invariance (i.e. coding for speed in spite of large changes in stimulus’ frequency range) would not explain the increased firing observed along the iso-scale axis (Fig 4d, red histogram; MI_135°; iso-scale_ > 0). This behavior can however be explained with a model in which PMLS neurons act as detectors of speed differences. Individual and compound motion clouds are first processed by a bank of spatio-temporal filters (Fig 4e; see *Materials and methods*), their outputs are then linearly combined to build selectivity for speed differences (Fig 4e). This model accurately captured the pattern of gain modulation observed in oriented units (Fig 4f; correlation between measured and predicted MI_*φ*_; Pearson’s r(304) = 0.47; p < 0.001). The receptive field estimated by the model was aligned along the bisector between the iso-speed and iso-SF line (Fig 4g, angle = 67.6 ± 7.8°), an orientation suggesting that *oriented* PMLS neurons respond preferentially to stimuli whose content varies both in spatial frequency and speed. Notably, this selectivity emerged later than orientation and spatial frequency, but earlier than temporal frequency and direction (Fig 4h), and highlight how PMLS neurons can be tuned for motion segmentation.

## Discussion

The experiments presented here were performed to explore the encoding, combination, and segmentation of naturalistic motion. Our results highlight that naturalistic motion is encoded in a temporally staggered manner, that nearby components are combined linearly, and that a a sub-population of neurons are equipped with receptive fields shaped for its segmentation by speed differences.

### Sequential selectivity to visual features

In contrast to simpler stimuli commonly used in experimental settings (Rust & Movshon, 2005), real world motion is not confined to a single point in Fourier space (Leon et al., 2012). We found that information content about the features of the stimulus could be decoded with different latencies, where orientation is decoded first, then spatial frequency, then temporal frequency, and finally direction. This progressing selectivity appeared not only during the first 100 ms of the stimulus (Smith et al., 2005), but also re-emerged with a specific oscillatory pattern (Dunn et al., 2022). This concomitant representation of information echoes information multiplexing (Panzeri et al., 2010) and highlights a mechanism for the representation of information in complex visual scenes. This time-evolving selectivity could be explained by the connections between PMLS and earlier visual areas (Jarosiewicz et al., 2012), and the prevalence of feed-forward and feedback mechanisms in the visual hierarchy (Celebrini et al., 1993).

### Linear combination of motion features in Fourier space

Our modeling efforts highlight that, when presented with compound motion clouds (i.e. combinations of motion clouds), neurons in area PMLS sum their responses to individual components. This behavior agrees with mechanisms at earlier stages of the visual system (Busse et al., 2009), supports feature pooling over Fourier space in motion processing (Masson & Castet, 2002), and echoes mechanisms highlighted in multisensory contexts (Deneve & Pouget, 2004; Gu et al., 2006; Fetsch et al., 2012). By construction, our compound motion clouds shared the same uncertainty; neurons’ linear response likely reflect the Bayes-optimal channel integration strategy (Zemel et al., 1998), here within a single modality. Neurons’ reliable response to naturalistic stimuli (Vinje & Gallant, 2000) could thus be explained by their rich spatio-temporal frequency content fused into a single representation, a fusion explaining the apparent cohesiveness of motion clouds and naturalistic stimuli.

The mechanisms we report in our study were observed over spatio-temporal frequencies lying close in Fourier space. However, motion processing does not only entail the combination of motion information but also its segmentation from background information (Braddick, 1993), namely its causal inference (Körding et al., 2007). This trade-off between integration and segmentation explains why similarities between low-levels features guide the behavior of the network from a cooperative *averaging* over features space (Mingolla et al., 1992; Lisberger & Ferrera, 1997; Lorenceau & Zago, 1999) to a competitive *winner-take-all* mode (for a review, see Masson & Perrinet, 2012). In estimating the speed of naturalistic motion, lateral inhibition strengthens with increasing distance of speed channels from the iso-speed line (Inagaki et al., 2016; Gekas et al., 2017; Meso et al., 2022), that is channels corresponding to visual inputs at different speeds. Ultimately, motion representation by a neuronal population, its normalization (Simoncini et al., 2012), and its interpretation by downstream areas must remain flexible to meet task-demands (Spering & Montagnini, 2011).

### Neural bases of motion segmentation in Fourier space

Interestingly, we found a sub-population of units of PMLS sensitive to speed differences. While in early areas, few units are truly speed-tuned (Perrone & Thiele, 2001; Priebe et al., 2006), this behavior spreads at later cortical stages (Priebe et al., 2003) where receptive fields are no longer separable in Fourier space. In contrast to speed-tuned receptive fields, the observed sub-population’s response did not increase along the iso-speed axis, but along an axis in-between the speed and spatial frequency axes. This sub-population was sensitive to differences orthogonal to the axis that bisects the iso-SF and iso-speed line, which is the principal component of speed-tuned units in area MT (Priebe et al., 2003). Their center-surround architecture suggest that they likely stem from speed-tuned units. Their behavior echoes antagonistic surround influence on speed coding in MT neurons (Allman et al., 1985; Bradley et al., 1998) and their velocity-discriminating nature is suited for depth inference from motion parallax. They have been hypothesized for distinguishing self-movement and eye movements from object movements (Nakayama & Loomis, 1974), and for segmenting objects based on their velocity profile (Braddick, 1993). Our results provide some evidence of their existence and further stress the role of extrastriate areas in segmenting objects based on how likely they share a *common fate* (Lorenceau, 2010).

## Conclusion

The ferret visual system shares some similarities with other animal species in its processing of visual motion. Our results here provide a first explanation as to how visual motion is encoded, combined, and segmented. In particular, we have shown that neurons in the posteromedial lateral suprasylvian sulcus (PMLS) encode features sequentially, with early responses to its orientation and spatial frequency properties before its temporal frequency and direction. These neurons also combine information linearly in Fourier space, building a unified representation of complex visual scenes. Lastly, some units provide the necessary receptive fields to perform motion and scene segmentation based on speed and spatial frequency differences. Overall, our work shows how information from moving visual scenes can be encoded, integrated, and accordingly parsed into elements that determines what the animal perceives.

## Materials and methods

### Visual stimulation

We presented two different sets of stimuli: motion clouds (MCs; Figs 1a–c; Leon et al., 2012; Vacher et al., 2018) and compound motion clouds (CMCs; Gekas et al., 2017; Meso et al., 2022). They were generated by applying an envelope *g* in Fourier space to constrain the spatio-temporal profile of a white noise image *I* where *g* is a 3-dimensional Gaussian-shaped function defined in log-frequency space and log-frequency time, and parametrized by its mode *µ* and bandwidth *B* (full width at half maximum) in Fourier space. The mode determines the critical spatial frequency (SF) and temporal frequency (TF) of the stimulus. The given image *I* at each spatial location (*x, y*) over time *t* was computed as follow:

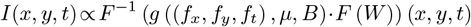

where *F* is the 3D Fourier transform, *F*^−1^ is the inverse Fourier transform, and *W* is a white noise image.

In the main experiment, the set of motion clouds included a central motion cloud whose critical spatial and temporal frequencies (SF_0_ = 0.07 cyc/deg; TF_0_ = 0.7 Hz; Figs 1a–c) were placed within the window of visibility of the ferret (see Li et al., 2006, for SF and TF; and Lempel & Nielsen, 2019, for speed) Around this central motion cloud, we chose 8 MCs that were at a 1 octave distance and arranged on a circle in log-frequency space and time (Fig 1d). The bandwidth *B* (^1^*/*_3_ octave) was chosen to minimize the marginal overlap between MCs in Fourier-space (here 5%). In a second experiment, the whole set of stimuli was shifted by 1 octave along the spatial frequency axis while maintaining the same speed (SF_0_ = 0.14 cyc/deg and TF_0_ = 1.4 Hz).

MCs and CMCs were generated in Matlab displayed on a gamma-corrected screen (Alienware AW2521HFA, 1920 *×* 1080, 60 Hz) placed 29 cm in front of the ferret eye, subtending an area of 86 *×* 55 degree of visual angle (dva), and centered on the ferret field of view. MCs and CMCs were scaled to ensure the same contrast (RMSE=34 cd/m^2^) across the stimuli set. They were vignetted by a 3 dva raised half-cosine window from black to gray. Their drifting direction was one of 8 equally spaced directions between 0 and 315°.

### Electrophysiological recordings

We recorded from the right hemisphere of two adult female ferrets (*Mustelia putorius furo*). Ferrets’ health and weights were closely monitored throughout the course of the experimental procedure. They were implanted with a stainless steel headpost at approximately 7 months old. They were sedated (medetomidine, 0.08 mg/kg, subcutaneous), and anesthetized (ketamine, 5 mg/kg, intramuscular; then 1–2% isoflurane) while their vitals (ECG, pulse, oxygenation, and rectal temperature) were continuously monitored. The animal’s skull was surgically exposed by an incision to the skin along the medial crest down to the neck and temporal muscles were removed from the medial crest to the beginning of the zygomatic arch. Stereotaxic coordinates of the posteromedial lateral suprasylvian sulcus (PMLS) were marked. A headpost was fixed to the midsagittal ridge, and a bone cement implant was molded to cover the skull. Ferrets were allowed a 2-week postoperative recovery during which they were administered antibiotics for 7 days and anti-inflammatory and analgesics for 4 days. They were then habituated to being head-restrained and underwent a second procedure. Their implant was drilled to build a well down to the skull, and a craniotomy was performed above the cortex. Across recording sessions, adjacent craniotomies were made to access the targeted region. At the end of the experiment, three marking penetrations were performed with a probe coated with a fluorescent lipophilic dye DiI (DiCarlo et al., 1996). The probe sat in place for 30 minutes before retraction to ensure sufficient dye transfer. After the final recordings, animals were deeply anesthetized with medetomidine and ketamine and euthanized with an overdose of pentobarbital. Their brain was immediately extracted and fixed in 4% paraformaldehyde in phosphate-buffered saline for at least 48 hours at 4°C. Brains were then coronally sectioned (50–100 µm thick) using a freezing microtome. Fluorescent labeling was imaged using a wide field epifluorescence microscope and electrode tracks were reconstructed by triangulating the DiI-marked penetrations relative to known skull landmarks and craniotomy coordinates. Nissl staining was performed on adjacent sections to verify cortical architecture and confirm localization within PMLS. Experiments were approved by the French Ministry of Agriculture (protocol authorization: 21022) and strictly complied with the European directives on the protection of animals used for scientific purposes (2010/63/EU).

For each recording, the ferret was placed in a custom-made tube inside a dark double-walled sound-proof (IAC) booth with the headpost secured to a holder. A linear probe (Assy77 M1, Cambridge Neurotech) maintained on a stereotaxic frame was descended through a puncture in the dura until finding responsive sites. Once responses had stabilized, stimulus presentation started for a about 1 hour long recording session. Raw traces were amplified, filtered, and digitized at a 30 kHz sampling rate (Siegle et al., 2017). After visual inspection of multi-unit activity and mechanical drift (Buccino et al., 2020), single units were extracted offline (Pachitariu et al., 2024). These so-obtained clusters were inspected, manually deleted, merged, or split, depending on their cross-correlograms, raw waveforms, and amplitude for a total of 530 units (experiment 1: 247 units for ferret A, and 62 units for ferret B; experiment 2: 177 units for ferret A, and 44 units for ferret B). Spikes were epoched from −200 ms to 700 ms around stimulus onset and binned in 1 ms time bins. For each unit, response latency was defined as the first bin after stimulus onset where average firing rate exceeded the spontaneous discharge rate by two standard deviations and followed by, at least, two consecutively increasing bins (Raiguel et al., 1999).

### Single-unit decoding analyses

The evoked activity of each neuron was filtered by a 5 ms gaussian smoothing kernel (Fig S1). To measure information content, the smoothed evoked activity for each unit underwent different decoding strategies:

a *full time-course* decoder (Figs 2, 4): the smoothed spike train during stimulus presentation was z-scored, denoised by PCA (keeping 90% of the variance, 18 PCs on average), and classified by a linear discriminant with time bins as features.

a *time-dependent sliding* decoder (Fig 2 and Fig S2a): the smoothed spike train inside a window of length *T* was z-scored, weighted by a ramp-up causal exponential window decaying from 1 to 0.05 and classified by a linear discriminant with *T* time bins as features. This procedure was repeated every 9 ms between −200 ms and +700 ms around stimulus onset, and for 8 windows’ durations *T* varying from 5 to 75 ms.

a *time-dependent integrating* decoder (Fig S2b): the smoothed spike train inside a window of length *T* was z-scored, denoised by PCA (keeping 90% of the variance), and classified by a linear discriminant with *T* time bins as features. The window truncated the spike train between −200 ms and +*T* ms around stimulus onset, with *T* increasing by 9 ms for every instance of the decoder.

a *cross-decoder* (Fig S3): the smoothed firing rate in a 10 ms sliding window was z-scored and underwent a linear discriminant analysis with time bins as features to find the direction maximally differentiating the conditions. The projection found this way was then applied to each time-steps before the window was moved by 2 ms.

These decoders were used on the 4 main attributes of the MCs analyzed, namely orientation, spatial frequency, temporal frequency, and direction. For spatial frequency (resp. temporal frequency), only the response to the MCs lying along the central line of equal temporal frequency (resp. spatial frequency) were used. For direction, each decoder was tested by grouping 3 adjacent orientations and testing against the 3 opposite orientations (the highest performance being retained). Classification accuracy was estimated by a 5-fold stratified cross-validation procedure (Varoquaux, 2018), and significance was assessed by deriving the chance distribution over 500 iterations of the decoder with randomly shuffled class labels, with a significance-level of 5% taken as threshold.

### Modeling responses to compound motion clouds

Responses to compound motion clouds (CMCs) were described by Poisson process whose rate followed a generalized non-linear summation model (defined by a p-norm):

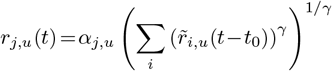

with *r*_*j,u*_ the response of unit u to the CMC *j* and 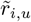 the smoothed evoked activity for motion cloud *I*_*i*_, namely

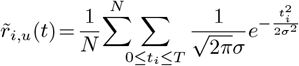

with *N* the number of trials, *t* taken every 1 ms, *T* = 500 ms the duration of the stimulus presentation, and *σ* the smoothing constant.

In this model, the parameters *α* and *γ* controlled the scaling and shape of the summation surface (for similar approach, see Britten & Heuer, 1999). Model’s behavior thus varied from a *summation* (*γ* = 1) to a *winner-take-all* regime (*γ* = ∞). For each unit, the *summation* (*γ* = 1), the *winner-take-all* (*γ* = ∞) and the *full model* (*γ* free) were fitted by maximum-likelihood, allowing the responses 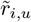 a temporal jitter *t*_0_ up to 10 ms around stimulus onset. Models’ goodness of fit were compared via the Akaike information criterion (AIC).

For the model that gave the relatively best description of the data at the level of the population, we modeled the scaling factor *α*_*j,u*_ as following a normal distribution centered on *λ*_*j*_ and whose variance was proportional to *λ*_*j*_ and inversely proportional to the firing rate of each unit *u*. Population’s parameters were compared to the predictions of a linear system (*λ*_*j*_ = ^1^*/*_2_ or ^1^*/*_3_ depending on the number of MCs).

Model’s performance was assessed by measuring, for each unit, the Pearson correlation between the predicted and measured firing rate. Since the amount of variance that can be explained by the best fitting model is limited by trial-to-trial variability rather than model miss-specifications, we compared these values obtained this way with a test-retest approach. For each unit, we computed the ceiling correlation (r_ceiling_), that is the Pearson’s correlation between the average response during CMC presentation and the smoothed firing rate for the same stimulus presentation in non-overlapping portions of the same recording in a 5-fold cross-validated manner (again allowing the underlying response a jitter up to 10 ms). For each CMC, we identified units with robust responses by deriving a chance distribution through 500 iterations of the same procedure with individual spikes corrupted by a uniformly distributed temporal jitter between −5 ms and +5 ms.

### Subpopulation analysis

In the second experiment, we also presented compound motion clouds (CMCs) composed of 3 individual MCs. These stimuli were built by combining MCs lying along the iso-temporal, iso-speed, iso-spatial, or iso-scale lines (Figs 4a, b). Thus, they all shared the same mean spatial and temporal frequency, and speed but their power varied along different angles *φ* in Fourier space. Notably, the CMCs aligned along the iso-speed (*φ* = 0°) and iso-scale (*φ* = 135°) axes shared the same marginal power along the spatial and temporal axis and only differed in their speed distribution.

The population of neurons was separated into two sub-populations depending on whether a full time-course decoder (see above) reached the 5% decoding significance level. For each unit *u*, and each axis in Fourier space *ϕ*, the modulation index MI_*φ,u*_ was defined as the relative firing rate increase when the power of the stimulus was increased along axis *φ*: MI_*φ,u*_ = (*R*_*φ,u*_ − *R*_0,*u*_)*/R*_0,*u*_ with *R*_*φ,u*_ (resp. *R*_0,*u*_) the average firing rate of unit *u* to the CMC aligned along *ϕ* (resp. central motion cloud) during the 100–500 ms post-stimulus onset interval.

### Population receptive field model

The observed modulation pattern was explained by a feed-forward computational model of motion cloud processing where PMLS neurons acted as speed difference filters. Their activity *m*_*φ,u*_ for each CMC aligned along axis *φ* (denoted *s*_*φ*_) was computed through a filter (here a difference between a circular and an oriented Gaussian) as follows:

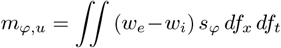

with *w*_*e*_ and *w*_*i*_ being 2-dimensional Gaussian functions in log-frequency space centered on 0, normalized to 1, shared across units, and parametrized by their covariances matrices 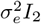 for *w*_*e*_, and 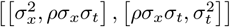 for *w*_*i*_. This model was fitted by maximum-likelihood to the firing rate *R*_*φ,u*_ of the units that decoded the iso-speed and iso-scale CMCs. The variability of its parameters was estimated by bootstrap resampling (1000 estimates).

### Eye tracking

Eye movements and pupil size were recorded with a Basler camera (Fig S5; acA8800, 64 Hz) in a control experiment without electrophysiological recordings. The viewing conditions and stimuli were the same as in the experiment but the ferret’s right eye was lit by a 840 nm infrared light. The stimuli sequence was preceded and followed by a calibration stimulus during which the screen was flashed from white (100 cd/m^2^) for 50 ms to black (0 cd/m^2^) for 450 ms. Pupil’s size was extracted offline by home-made routines. These data were epoched from −200 ms to 700 ms around stimulus onset. Trials with blinks were discarded and the remaining trials were centered and averaged across stimuli presentations.

## Acknowledgments

We thank Jonatan Nordmark for help with the initial stage of the recordings, Jonathan Vacher for mathematical inputs on stimulus construction and decoding, Samuel Garcia and Rémi Proville for spike-sorting recommendations, Flavien Feral for help with setting-up the eye-tracking device, Thibaut Doisy for animal care, David Hing and the Pasteur histology platform for histology, Najib Majaj for suggestions and experimental recommendations, and Guillaume Masson and Peter Neri for careful reading of the manuscript and precious comments. This work was supported by a ED3C (École Doctorale Cerveau, cognition, comportement) PhD fellowship to TS, the Institut Universitaire de France and ANR-19-CE37-0016 (Agence Nationale de la Recherche) to YB, ANR-19-NEUC-0003-01 to PM, and ANR-17-EURE-0017, ANR-10-IDEX-0001-02, and PSL (Université Paris Sciences & Lettres).

## Competing interests

The authors declare no competing interests.

## Supplementary material

**Fig S1.**
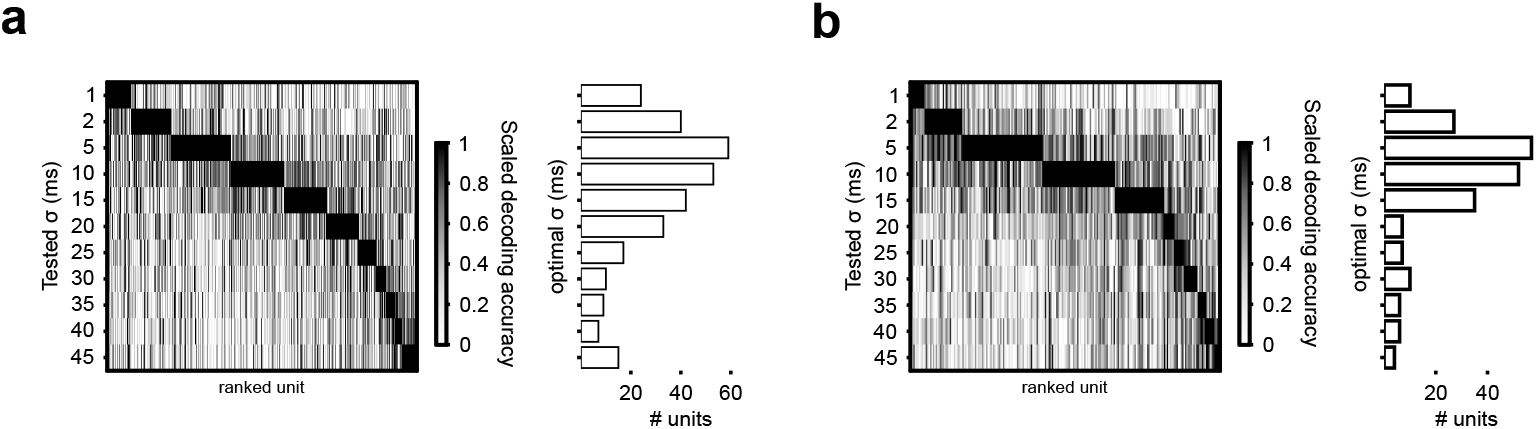
Grid-search for the smoothing parameter. For each unit, the firing rate time-course was smoothed with a given constant *σ* before undergoing decoding. Histogram on the right report the distribution of the optimal *σ* for experiment 1 in **(a)** and experiment 2 in **(b)**.

**Fig S2.**
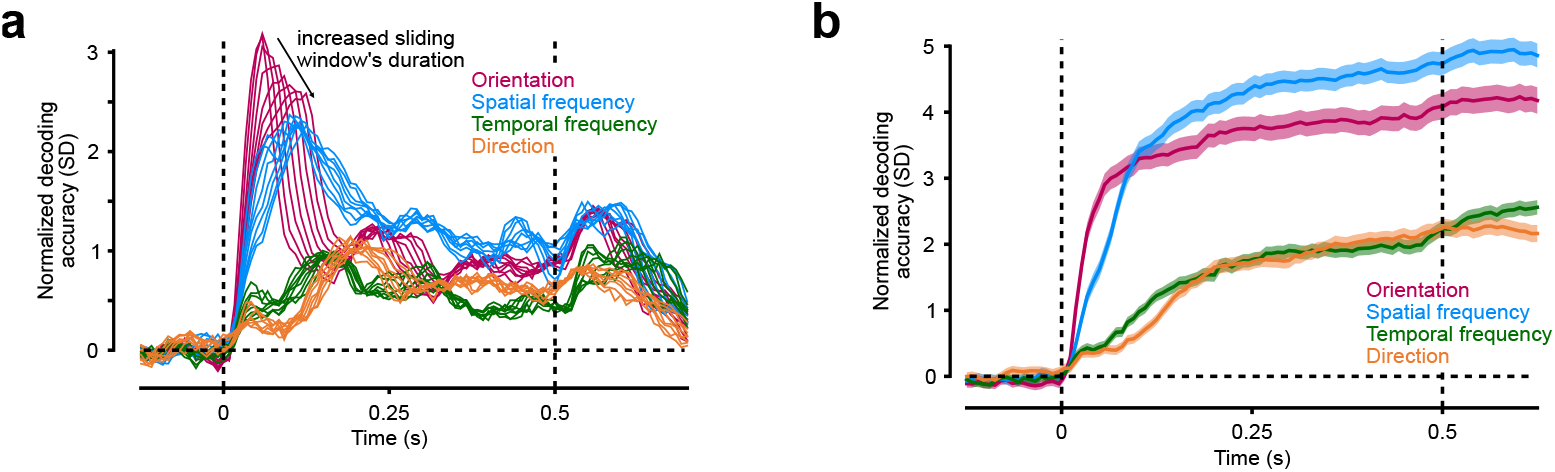
Decoding with varying parameters. **(a)** Decoding of the four main features for different sliding windows sizes ranging from 5 ms to 75 ms. **(b)** Decoding of the four main features with an integrating window (see *Materials and methods*).

**Fig S3.**
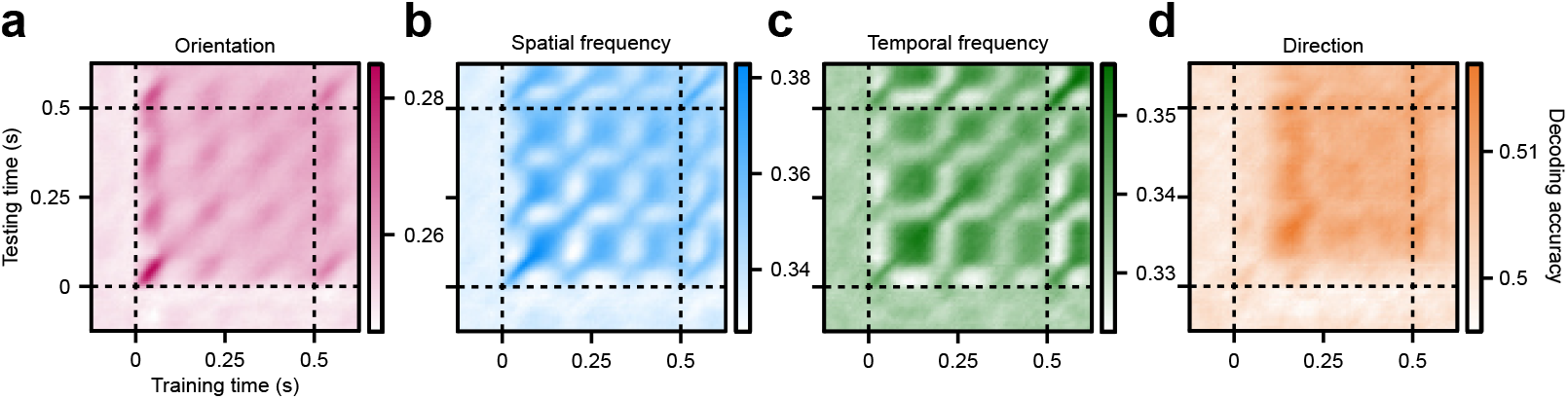
Cross-decoding analysis. For each features of the stimulus **(a)** Orientation, **(b)** Spatial frequency, **(c)** Temporal frequency, and **(d)** Direction the time-course of the encoding of features encoding was tested.

**Fig S4.**
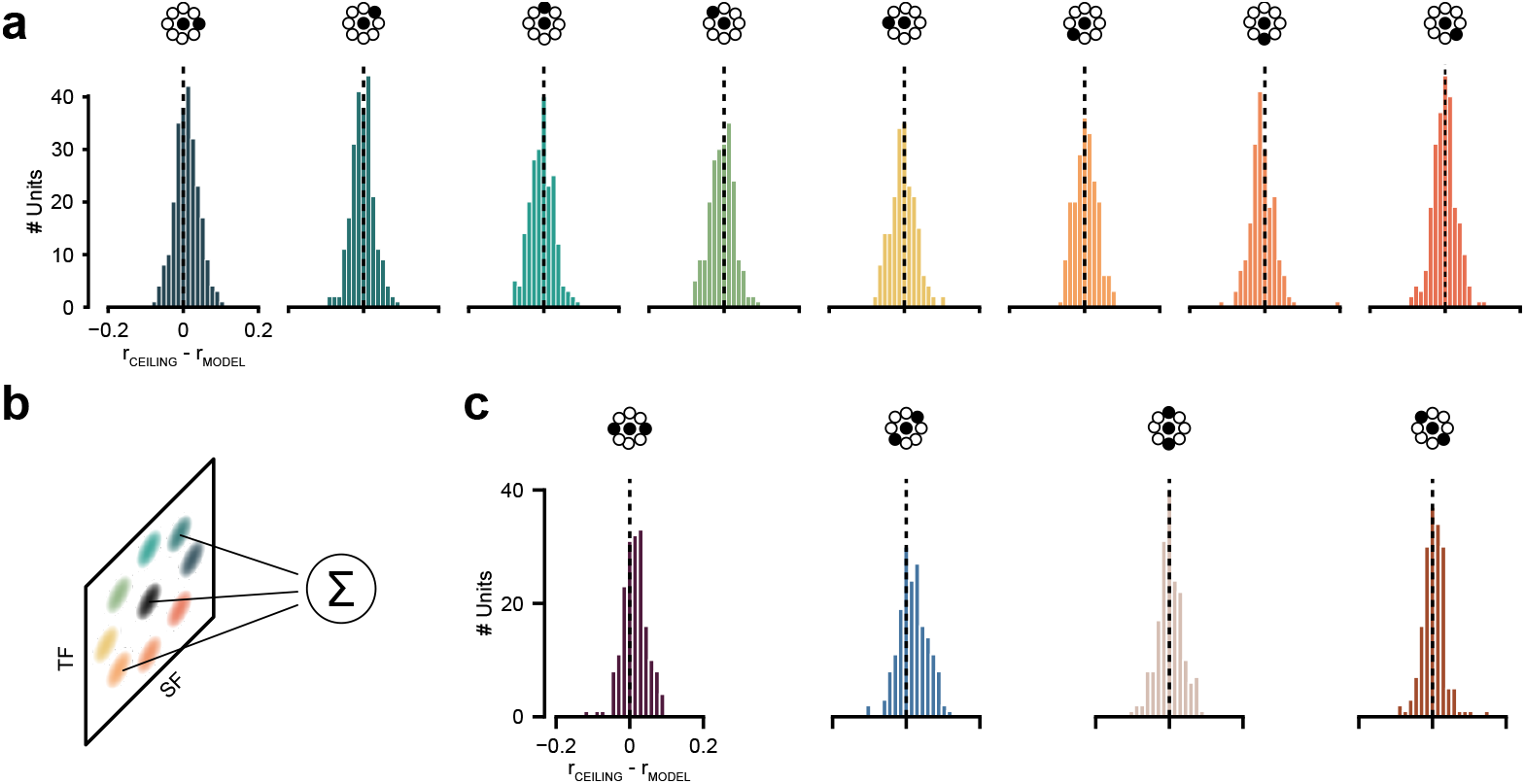
Fit for each individual motion clouds. **(a)** Difference between the correlation of the test-retest model (r_ceiling_) and the correlation of the linear model (r_model_) for each of the individual motion clouds. **(b)** Illustration of the extension of the model to the triplets of motion clouds (see Fig 4). **(c)** Difference between the correlation of the test-retest model and the correlation of the linear model for the four compound motion clouds triplets.

**Fig S5.**
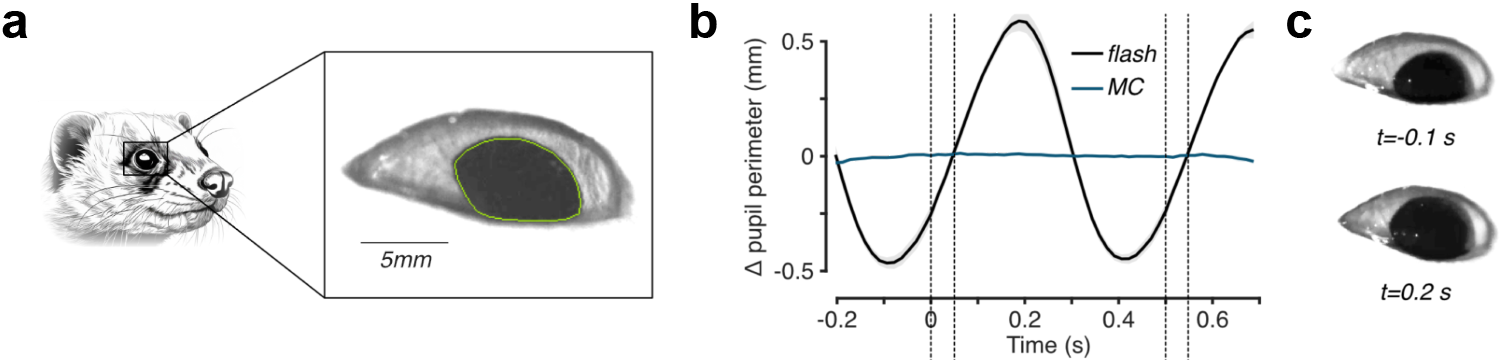
Pupil segmentation. **(a)** Example of the segmentation of the pupil during the set of control experiments. **(b)** Pupil size response for the motion clouds in blue and the control stimulus i.e. the screen flashing from black to white. The onset and offset of this flash stimulus are indicated by the vertical dashed lines. **(c)** Example raw pupil image at 0.1 s pre-flash onset and 0.2 s post-flash onset.

## Notes

### Competing Interest Statement

The authors have declared no competing interest.

